# Discovery of repurposing drug candidates for the treatment of diseases caused by pathogenic free-living amoebae

**DOI:** 10.1101/2020.05.13.093922

**Authors:** Christopher A. Rice, Beatrice L. Colon, Emily Chen, Mitchell V. Hull, Dennis E. Kyle

**Affiliations:** Department of Cellular Biology, University of Georgia, Athens, Georgia, USA; Center for Tropical and Emerging Global Diseases, University of Georgia, Athens, GA, USA, 30602; Department of Infectious Diseases, University of Georgia, Athens, Georgia, USA; Calibr at Scripps Research, La Jolla, CA, USA, 92037

**Author notes:** Current address: Biological Chemistry and Drug Discovery Department, University of Dundee, Dundee, Scotland, United Kingdom, DD1 5EH (Beatrice L. Colon). Corresponding authors (DEK); (CAR).

**Keywords:** Phenotypic screening, drug discovery, *Naegleria fowleri*, primary amoebic meningoencephalitis, *Acanthamoeba castellanii*, granulomatous amoebic encephalitis, *Acanthamoeba* keratitis, cutaneous amoebiasis, *Balamuthia mandrillaris*, *Balamuthia* amoebic encephalitis, antiparasitic agents, ReFRAME drug library, Calibr-Scripps

## Abstract

Diseases caused by pathogenic free-living amoebae include primary amoebic meningoencephalitis (*Naegleria fowleri*), granulomatous amoebic encephalitis (*Acanthamoeba* spp.), *Acanthamoeba* keratitis, and *Balamuthia* amoebic encephalitis (*Balamuthia mandrillaris*). Each of these are difficult to treat and have high morbidity and mortality rates due to lack of effective therapeutics. In pursuit of repurposing drugs for chemotherapies, we conducted a high throughput phenotypic screen of 12,000 compounds from the Calibr ReFRAME library. We discovered a total of 58 potent inhibitors (IC_50_ <1 μM) against *N. fowleri* (n=19), *A. castellanii* (n=12), and *B. mandrillaris* (n=27) plus an additional 90 micromolar inhibitors. Of these, 113 inhibitors have never been reported to have activity against *Naegleria*, *Acanthamoeba* or *Balamuthia*. Rapid onset of action is important for new anti-amoeba drugs and we identified 19 compounds that inhibit *N. fowleri in vitro* within 24 hours (halofuginone, NVP-HSP990, fumagillin, bardoxolone, belaronib, and BPH-942, solithromycin, nitracrine, quisinostat, pabinostat, pracinostat, dacinostat, fimepinostat, sanguinarium, radicicol, acriflavine, REP3132, BC-3205 and PF-4287881). These compounds inhibit *N. fowleri in vitro* faster than any of the drugs currently used for chemotherapy. The results of these studies demonstrate the utility of phenotypic screens for discovery of new drugs for pathogenic free-living amoebae, including *Acanthamoeba* for the first time. Given that many of the repurposed drugs have known mechanisms of action, these compounds can be used to validate new targets for structure-based drug design.

**Author Summary:** Free-living amoebae (FLA) are ubiquitous in soil and freshwater and most are non-pathogenic to people; however, three different pathogenic FLA have been found to cause severe, most often fatal diseases in humans. Due to poor detection and inadequate treatment options available for pathogenic FLA, the fatality rates are still > 90% for the diseases caused by *Balamuthia mandrillaris*, *Naegleria fowleri*, and *Acanthamoeba* spp. With hundreds of cases in the United States and many more cases reported worldwide, there is still an urgent clinical need for effective diagnosis and specific treatments discovered against these opportunistic parasites. Drug repurposing is a powerful approach for drug-discovery because it significantly improves the discovery time, reduces the amount of resources, and decreases costs required to advance lead candidate drugs of interest into the clinic. This is extremely helpful for neglected diseases including pathogenic FLA where there is a need for new active therapies with limited budgets. This report addresses the discovery of new active drugs with potential for repurposing, multiple new drug classes that inhibit pathogenic FLA, and numerous putative drug targets that can be used as tools for further investigation and structure-based drug design.

## Introduction

*Naegleria fowleri*, the causative agent of primary amoebic meningoencephalitis (PAM), was first discovered to be a human pathogen by Fowler and Carter after four fatal infections from 1961-1965 in Adelaide, Australia^1^. From 1962-2018, 145 cases have been reported in the United States (US)^2^, with several hundreds of cases recorded around the world^3–7^. *Naegleria*, being thermophilic, thrives in warm fresh waters where it can exist as one of three forms: the trophozoite form (active feeding stage), flagellated form (motile swimming stage), or as a cyst form (protective dormant stage). The trophozoite is thought to be the only infective stage of *N. fowleri*. PAM is not a notifiable disease in the US, and due to common misdiagnosis as bacterial or viral meningitis, the number of reported cases is likely to be a significant underestimate of disease burden globally. PAM is commonly associated with recreational water activities in which water is forcefully inhaled into the nasal cavity. For example, swimming, jumping into water without holding the nose or wearing a nose clip, water sports (wake boarding, water skiing, jet skiing, white water rafting), and more recently increased use of neti pots for sinus/nasal irrigation or religious practice of nasal cleansing, are all risk factors for PAM^8–11^. Once *N. fowleri* enters the nasal cavity, amoebae traverse the cribriform plate the olfactory nerve to reach the frontal cerebral cortex of the brain where they cause hemorrhagic meningoencephalitis with the classical PAM symptoms of severe headache, stiff neck, hallucinations, seizures, coma, and, in 97% of cases, death^12,13^. With gradually increasing awareness and better diagnostic tools (real-time PCR, LAMP assays, antigen detection and serological testing) more cases are being diagnosed faster. Key to the few instances of PAM survival is early diagnosis and a combinational treatment regimen that includes amphotericin B, azithromycin, fluconazole, miltefosine, rifampin and dexamethasone. Miltefosine, a breast cancer chemotherapeutic agent and anti-leishmanial drug, was repurposed for treatment of FLA diseases after the discovery of activity in *in vitro* and *in vivo* models^14,15,47^. Several recent PAM survival cases included inducement of therapeutic hypothermia (cooling of the core body temperature to 32-34°C) to better manage intracranial pressure; this technique also reduces reactive oxygen and nitrogen species, neural apoptosis, and proinflammatory cytokine levels, all of which increase brain injury through hyperinflammation^16,17^.

*Acanthamoeba*, unlike *Naegleria*, causes disease in immunocompromised and immunocompetent individuals; the first human infections were described in 1958 as the cause of granulomatous amoebic encephalitis (GAE)^18^. To date, 22 different sequence types of *Acanthamoeba* have been reported by the community based on >4% pairwise sequence difference in the *Acanthamoeba* specific amplifier (ASA) within the 18S rRNA gene sequence^19,20^. Various genotypes have been more commonly associated with different infections of *Acanthamoeba* disease; for example, T4 is the most commonly isolated genotype and primarily associated with GAE and *Acanthamoeba* keratitis, whereas genotypes T1, T10 and T12 are associated with GAE. Genotypes T4, T3 and T11 are more commonly found in *Acanthamoeba* keratitis infections and more recently genotypes T5 and T17 have been associated with cutaneous amoebiasis^21,22^. One of the major problems and concerns with all *Acanthamoeba* infections are that parasites can encyst into their protective dormant cyst stage and resist treatment within all of the tissues it can infect^22–24^. Treatment is terminated when the patient’s symptoms improve, however persistent parasites can excyst and cause further pathology and disease. Recrudescence has been reported up to 5 years after initial infection and treatment^25,26^.

*Acanthamoeba* keratitis (AK) is an excruciating and debilitating infection of the cornea. It is the most common disease caused by *Acanthamoeba* species, mainly in association with improper contact lens wearing, although non-contact lens wearing AK cases have been reported^27^. Diagnosis is challenging and, if not treated aggressively, visual impairment or blindness can occur^28^. Moderately successful treatment, if applied early, has been achieved by using the combinational chemotherapy of 0.02-0.2% chlorhexidine gluconate, 0.02-0.06% polyhexamethylene biguanide (PHMB), and 0.1% propamidine, with or without neomycin, 1% azoles, or other amidines (e.g., 0.1% hexamidine)^29,30^. Depending on the clinical response, chemotherapy with topical agents can vary from months to years. Unfortunately, most of the current chemotherapy agents are cytotoxic to corneal cells and, as an example, at one eye care clinic, approximately 25% of confirmed AK patients required therapeutic keratoplasty (corneal transplantation) or in extreme cases enucleation of the eye^31^.

Cutaneous amoebiasis lesions and nasopharyngeal infections caused by various *Acanthamoeba* species are mainly in immunosuppressed patients such as human immunodeficiency virus (HIV) infections, chemotherapy recipients, those with immunodeficiency syndromes, and transplant recipients^32^. Only 5 cutaneous cases have been reported in immunocompetent patients with a unique case of oste-cutaneous acanthamoebiasis (bone involvement) due to traumatic inoculation through an injury sustained playing sports^33^. All other cases are through *Acanthamoeba* parasites infecting through cuts or breaks in the skin or via the nasal passage. With a case mortality of ~70% in the absence of GAE and almost always fatal with central nervous system (CNS) involvement, better treatments for successful patient outcomes need to be investigated^24^. There is no clinical agreement or treatment recommendation for cutaneous lesions; rather healthcare providers have depended upon research to decide which agents to use for treating *Acanthamoeba* infections. Broadly, cutaneous cases have been successfully treated with combinations of amphotericin B, pentamidine isethionate, chlorhexidine gluconate, 5-fluorocytosine, fluconazole, itraconazole, ketoconazole, azithromycin, clarithromycin sulfadiazine, and pyrimethamine, along with the debridement of sinuses in cases of sinusitis^34–36^.

From skin or lung infections, *Acanthamoeba* can spread to the CNS through haematogenous dissemination where it can attach, and break down the blood-brain barrier, then infect brain tissue causing granulomatous amoebic encephalitis (GAE)^24^. Similarly to PAM, GAE is most commonly diagnosed post mortem, since the clinical symptoms of headache, vomiting, nausea, confusion, fever, ataxia, increased cranial pressure, and seizures that resemble meningitis caused by bacteria or viruses leading to misdiagnosis^37^. GAE is found in patients with preexisting underlying diseases. It has a slower more chronic disease progression of the course of several weeks to months before clinical symptoms arise and an accurate diagnosis can be made. GAE has a > 90% mortality rate. Currently, a small number of successful outcomes have resulted from treatment with a mixture of combinational chemotherapies including antifungal azoles (clotrimazole, miconazole, ketoconazole, fluconazole, itraconazole, or voriconazole), antifungals (amphotericin B, flucytosine, caspofungin, pentamidine isethionate, hydroxystilbamidine, 5-fluorocytosine), antibiotics (azithromycin, rifampin, trimethoprim/sulfamethoxazole, sulfadiazine, chloramphenicol, paromomycin, polymyxin) and more recently miltefosine^24,38^.

*Balamuthia mandrillaris* has a similar pathobiology to *Acanthamoeba* and is the causative agent of the brain disease *Balamuthia* amoebic encephalitis (BAE); it has also been reported to cause similar cutaneous amoebiasis. Clinical cases of BAE have been identified with or without cutaneous skin involvement^39^. Since the discovery of *B. mandrillaris* as an etiological agent in 1986, 108 BAE cases have currently been reported by the CDC between 1974-2016; the clinical cases described in the years prior to 1990 were retrospectively diagnosed^40,41^. Similarly to the other amoebae, the treatment of *B. mandrillaris* diseases has ultimately revolved around success in a small number of surviving cases and limited *in vitro* drug susceptibility data. With only 9 successfully treated BAE cases in the US, descriptions of successful combinational chemotherapies include antifungal azoles (ketoconazole, fluconazole, or voriconazole), antifungals (amphotericin B, flucytosine or pentamidine isethionate), antibiotics (azithromycin, ceftriaxone, clarithromycin, ethambutol, isoniazid, rifampin, trimethoprim/sulfamethoxazole, sulfadiazine, pyrazinamide, doxycycline, metronidazole, or minocycline), antiparasitics (albendazole or miltefosine), steroids (prednisolone, dexamethasone, clobetasol propionate), antipsychotics (trifluoperazine, thioridizine) and antivirals (acyclovir)^41^. The most recent survivor in 2014 was treated in combination with albendazole, liposomal amphotericin B, azithromycin, clarithromycin, fluconazole, flucytosine, sulfadiazine and miltefosine, highlighting the fact that some of the drugs described in the aforementioned combinational cocktail were used for suspected and misdiagnosed bacterial or viral meningitis. To further complicate treatments, both the infective and destructive trophozoite stage and the dormant cyst stage of *B. mandrillaris* have been identified during infections^42^.

These high mortality rates indicate that there is a significant unmet medical need to discover new drugs that are efficacious against *Naegleria*, *Acanthamoeba* and *Balamuthia* trophozoites and/or cysts and have increased selectivity to the pathogen over infected dead-end host. Given the lack of major pharmaceutical company involvement in discovery of new drugs for the diseases caused by amoebae, drug repurposing is an appealing approach to overcome the unmet medical needs.

The California Institute for Biomedical Research (Calibr), <u>Re</u>purposing, <u>F</u>ocused <u>R</u>escue, and <u>A</u>ccelerated <u>Me</u>dchem (ReFRAME) drug repurposing library is an open access collection of ~12,000 commercially available or synthesized small molecules that are best-in-class, chemically diverse compounds with known safety profiles of drugs that have previously been tested in humans and animals. The library is comprised of roughly 38% FDA approved drugs, 30% Phase II/III clinical trial candidate drugs, 19% Phase I/0 lead drugs, 3% preclinical developmental compounds, and ~10% of compounds in an undetermined clinical stage. About 80% of the ReFRAME collection was assembled by firstly extensive patent mining of small molecules with known human safety profiles and then filtering candidate molecules through three widely used commercial drug competitive pharma research and development intelligence databases (Clarivate Integrity, GVK Excelra GoStar, and Citeline Pharmaprojects)^43^. The ReFRAME drug repurposing library was assembled to provide non-profit and academic researchers access to a high-valued, chemically diverse, and large compound library to enable rapid screening of compounds for efficacy against a plethora of diseases. All screening data and reconfirmed hits are deposited into a publicly accessible portal to allow for open mining of activity data (https://reframedb.org); this to accelerate drug development for new therapeutics and off-label indications, often for rare or neglected diseases with unmet clinical needs.

The goal of this study was to identify clinically approved compounds that have never been reported to have activity against pathogenic amoebae so they can potentially be repurposed for the treatment of several amoebic diseases. We screened the Calibr ReFRAME library and identified 58 nano- and 90 micro-molar inhibitors against *N. fowleri*, *A. castellanii* and *B. mandrillaris*; 113 of which have never been described to be active against *Naegleria*, *Acanthamoeba* or *Balamuthia*. In addition, we identified 19 compounds that inhibit *N. fowleri in vitro* within 24 hours. These compounds inhibit *N. fowleri* quicker than the standard drug regimen that is currently recommended by the CDC for the treatment of PAM. With nearly the entire ReFRAME library reaching preclinical safety profiling or clinical development this makes the library a very attractive and valuable set of compounds for the exploration of off-label therapeutic indications from those originally targeted diseases, especially for the treatment of pathogenic FLA.

## Materials and methods

### Amoeba culture

The *Naegleria fowleri* isolate (ATCC 30215) used in these studies was originally isolated from a 9-year old boy in Adelaide, Australia, that died of PAM in 1969. This isolate was purchased from the American Type Culture Collection (ATCC). *N. fowleri* trophozoites were grown axenically at 34°C in Nelson’s complete medium (NCM). The *Acanthamoeba castellanii* T4 isolate (ATCC 50370) used in these studies was isolated from the eye of a patient in New York, NY, in 1978. This isolate was also purchased from ATCC. *A. castellanii* trophozoites were grown axenically at 27°C in Protease Peptone-Glucose Media (PG). The *Balamuthia mandrillaris* isolate (CDC: V039; ATCC 50209), used in these studies was originally isolated from a pregnant Baboon that died of BAE in 1986 at San Diego Zoo, San Diego, USA. This isolate was kindly donated by Dr. Luis Fernando Lares-Jiménez ITSON, Mexico. *B. mandrillaris* trophozoites were grown axenically at 37°C in BMI medium. All reagents for media were purchased from Sigma-Aldrich (St. Louis, MO). *Naegleria*, *Acanthamoeba* and *Balamuthia* trophozoites from the logarithmic phase of growth were used for all drug susceptibility screening and secondary screening assays.

### *In vitro* drug susceptibility assays

The phenotypic drug susceptibility assays were performed as previously described using CellTiter-Glo v2.0 (CTG) (Promega, Madison, WI)^44,49,63^. The ReFrame collection of 12,000 compounds assembled at Calibr–Scripps were prepared in 384-well (Corning, White Flat bottom 3570) plate format at screening concentrations of 5μM (30nL of 10mM stock concentration/well in dimethylsulfoxide (DMSO)) using the Labcyte Echo 555 for acoustic compound dispensing. The first round was to screen in a single-point assay with 3,000 *Naegleria*/well, 600 *Acanthamoeba*/well or 4,000 *Balamuthia*/well in a total volume of 60μl. Prior to initiating drug screening, a maximum of 2% total hits from the library were predetermined for follow up from Calibr-Scripps for all screening projects. Drug spotted plates from Calibr-Scripps were left, at room temperature, to thaw overnight and parasites were plated using the Biomek NX^p^ automated liquid handler (Beckman Coulter). Hits were defined as compounds that produced > −40% (for *Naegleria*) or > −50% (for *Acanthamoeba* and *Balamuthia*) of normalized growth inhibition compared to the positive growth control (0.5% DMSO) and inhibitor controls that produced ~100% inhibition of amoebae cell populations. Posaconazole, azithromycin and DB2385A all at 1μM were used for controls for *Naegleria* and *Acanthamoeba*; artovastatin, fluvastatin and simvastatin all at 1μM were used as controls for *Balamuthia*. The second round was to verify the initial active hits from round I, hits were identified and serially diluted in duplicate from 5μM to 2nM in a 1:3, 8 point dilution series in 384-well plates with the same respected number of amoebae for *Naegleria*, *Acanthamoeba* and *Balamuthia* with the exact same controls and concentrations for each respected amoebae to generate the concentration for half-maximal activity derived from the hill equation model (q*AC*_50_). Dose-response curve fitting was analysed with Genedata Analyser software using the Smart Fit function. All assayed plates were incubated at the genus optimum growth temperature described above and tested for a period of 72 hours.

### Rate of action assay

RealTime-Glo MT Cell Viability Assay (RTG; Promega, Madison, WI) previously described by Colon *et al.*, was used to assess the speed of inhibition of active compounds against *N. fowleri*^49^. Compounds were plated at 1x and 5x their respected IC_50_’s with 3 replicates per plate. The 96-well plates were prepared with 10μl of drug, 4,000 cells in 40μl, and 50μl of RTG reagent (enzyme and substrate mixture). The negative growth control was cells in complete media with RTG reagent; the positive growth control was complete media and RTG reagent alone. Plates were incubated at 34°C for 72 hours within a SpectraMax i3x plate reader (Molecular Devices; Sunnyvale, CA). Relative luminescence units were recorded kinetically every hour for 72 hours and analysed using Prism 8 (GraphPad Software; La Jolla, CA).

### *Acanthamoeba* cysticidal assay

We screened all of the reconfrimed hits for *A. castellanii* for cysticidal activity using a modified method previously published by Rosales *et al.*,^64^. The trophozoites of *Acanthamoeba castellanii* were encysted in Neff’s Encystment Media (NEM)^65^, (20 mM Tris-HCL [pH 8.8], 100 mM KCL, 8 mM MgSO4, 0.4 mM CaCL2, 1 mM NaHCO3) (all reagents were from Sigma-Aldrich, St. Louis, MO) at 37°C for 3 days. Cysts were harvested using a cell scraper and washed with 0.5% sodium dodecyl sulphate (SDS) to remove any remaining trophozoites and immature cysts. All compounds were diluted in NEM and serially diluted in 2-fold dilutions 6 times to yield a working stock concentration range of 100μM to 3.125μM with a final concentration of 1% DMSO. Ninety (90)μl of a stock concentration of 1.1 × 10^4^/ml cysts and 10μl of experimental compounds from the stock dilution plates were combined for screening into clear 96 well treated tissue culture plate (Corning, NY) for 24 hours at 27°C. All wells contained 100μl. At the 24 hour time point plates were centrifuged at 4000 rpm at 25°C for 10 minutes and washed with phosphate-buffered saline (PBS), twice. All wells except the positive cyst control (cysts in non-drug treated NEM, 100μl) received 100μl of PG growth medium. The plates were left to recover at 27°C for 120 hours (5 days). The plates were microscopically analysed for the excystation process and the presence of trophozoites was used as an indication of cyst viability.

### Statistical analysis

We used the Z’ factor as a statistical measurement to assess the robustness of our high-throughput screening assays (Sup Figures 1, 2 and 3). This factor uses the mean and standard deviation values of the positive and negative controls. The robust RZ’ factor replaces plate means to robust medians and standard deviations to robust standard deviations which identify true hits. These 2% predetermined library hits were then followed up with dose-response for hit validation. For the RealTime-Glo analysis we measured <u>a</u>rea <u>u</u>nder the <u>c</u>urve (AUC) (Sup figure X & Y) and used Multiple T tests, One unpaired T test, False Discovery Rate (FDR = 1%) using method: Two-stage step-up method of Benjamini, Kreiger and Yekutieli to identify significance. Significance is recorded as *P*= < 0.05 (*), 0.0021 (**), 0.0002 (***), < 0.0001 (****), > 0.05 (NS – non-significant).

## Results

### Phenotypic screening of the Calibr ReFRAME library

Our first round of screening was performed at a single dose of 5 μM against *N. fowleri*, *A. castellanii* and *B. mandrillaris* logarithmic trophozoites. We initially identified 230 hits (1.9% of the total library) that produced −40% normalized percent growth inhibition or better for *N. fowleri*, 184 hits (1.5% of the total library) that produced −50% normalized percent growth inhibition or better for *A. castellanii* and 220 hits (1.8% of the total library) that produced −50% normalized percent growth inhibition or better for *B. mandrillaris* (Fig. 1). Hits from the first round were confirmed by quantitative dose response assays (N=2); from these data we confirmed a total of 53 bioactive (0.44% of the total library) molecules from 10 drug classes that produced replicative inhibitory growth curves of *N. fowleri* (Table 1). From the *Acanthamoeba* screen, we reconfirmed 32 bioactive (0.26% of the total library) molecules from 8 drug classes that produced replicative inhibitory growth curves of *A. castellanii* (Table 2). We further identified 63 bioactive (0.52% of the total library) molecules from 15 drug classes that produced replicative inhibitory growth curves of *B. mandrillaris* (Table 3).

**Figure 1.**
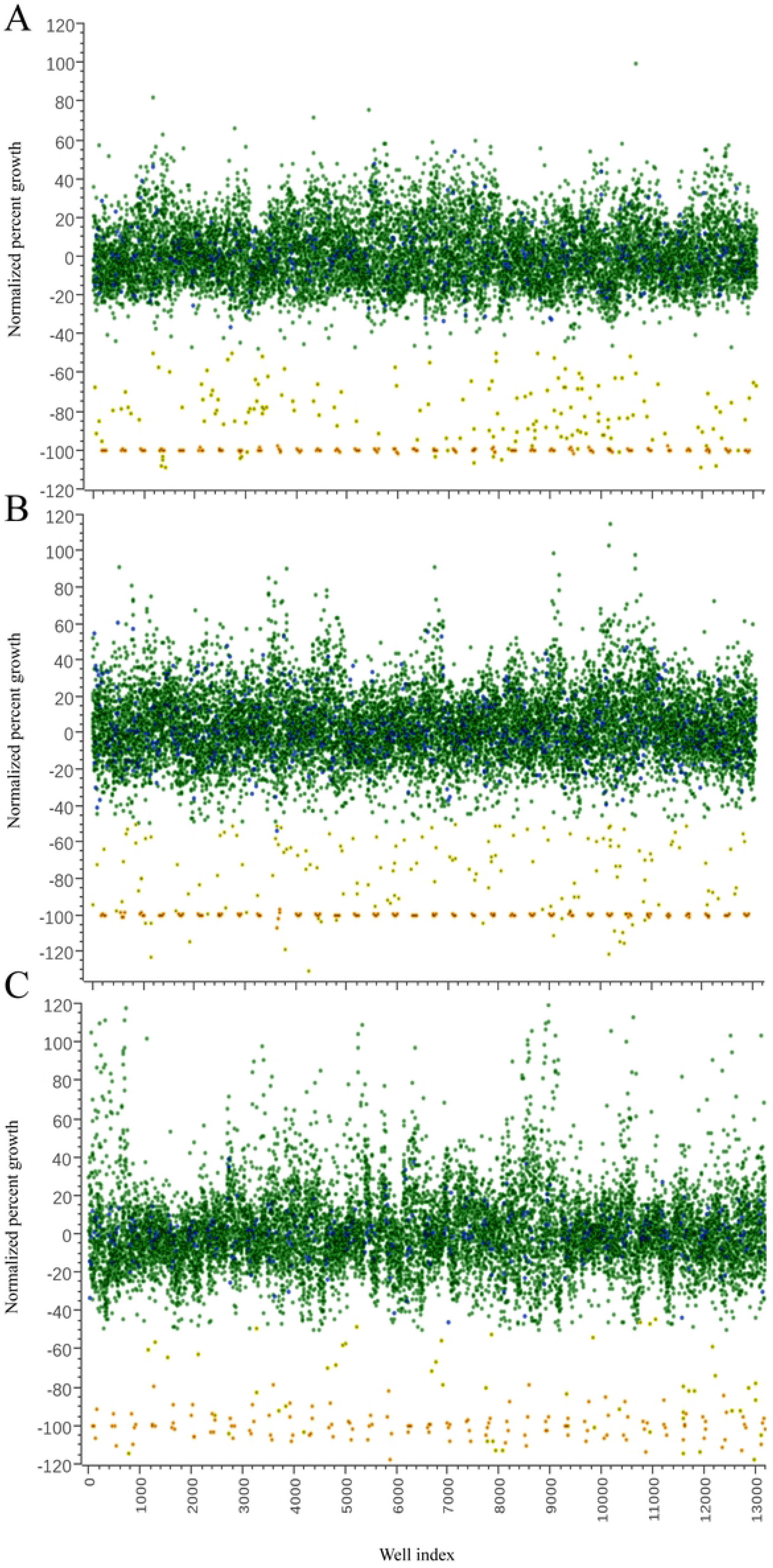
Scatter plot analysis of individual drug responses of the Calibr ReFRAME library at 5 μM. Single point scatter plot of the Calibr ReFRAME library screened at 5 μM against pathogenic A) *N. fowleri*, B) *A. castellanii* and C) *B. mandrillaris.* Each green circle represents individual compounds screened, the blue circles represent 0.5 % DMSO control (negative controls), the yellow circles represent the number of hits identified per well, the orange circles represent the median drug control (posaconazole for *Naegleria* and *Acanthamoeba*, and atorvastatin for *Balamuthia*) (positive controls). All values were normalized to percent growth per well. We used the RZ’ factor as a statistical measurement to assess the robustness of our high-throughput screening assays (Sup. Fig. 1).

**Table 1:**
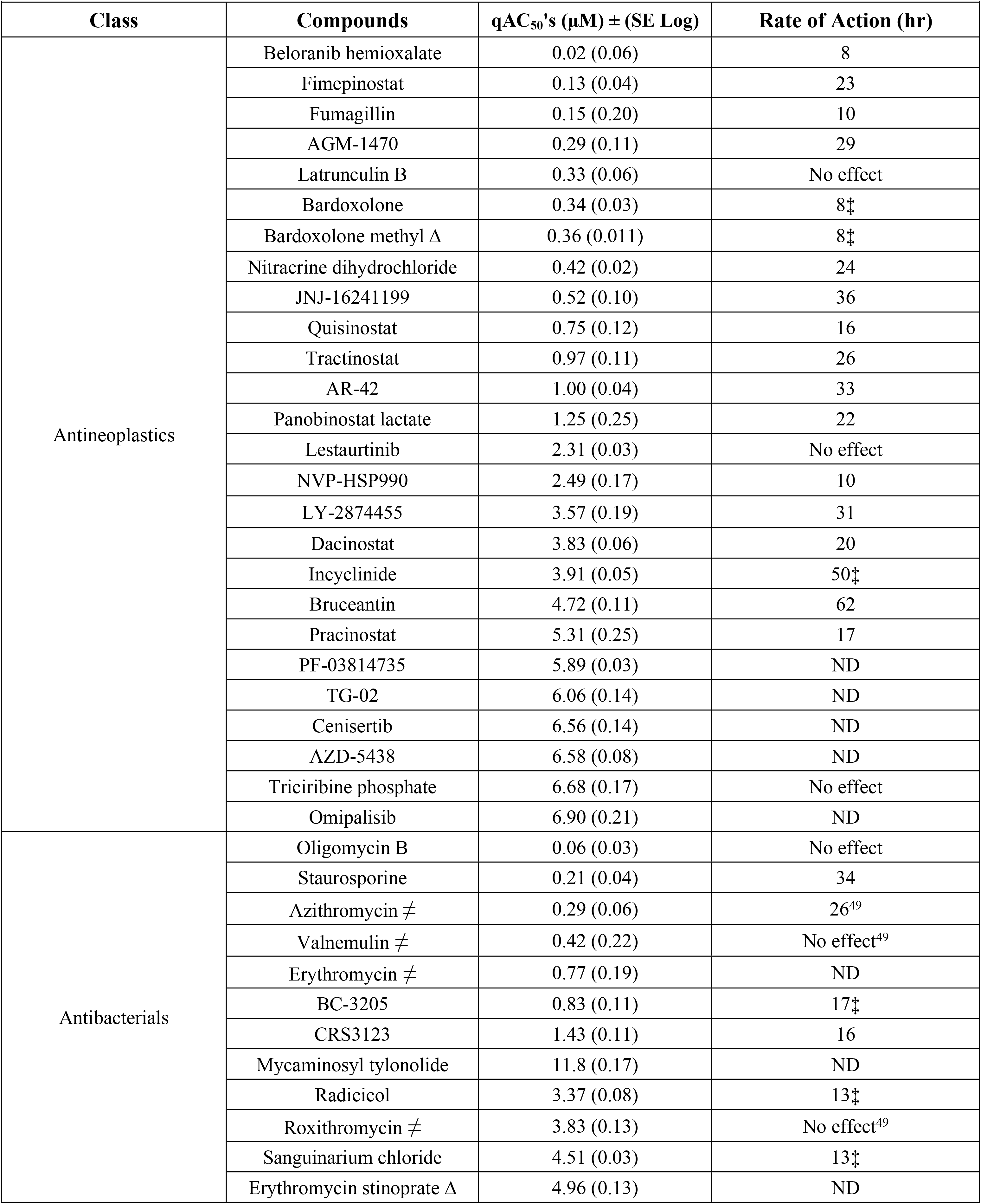
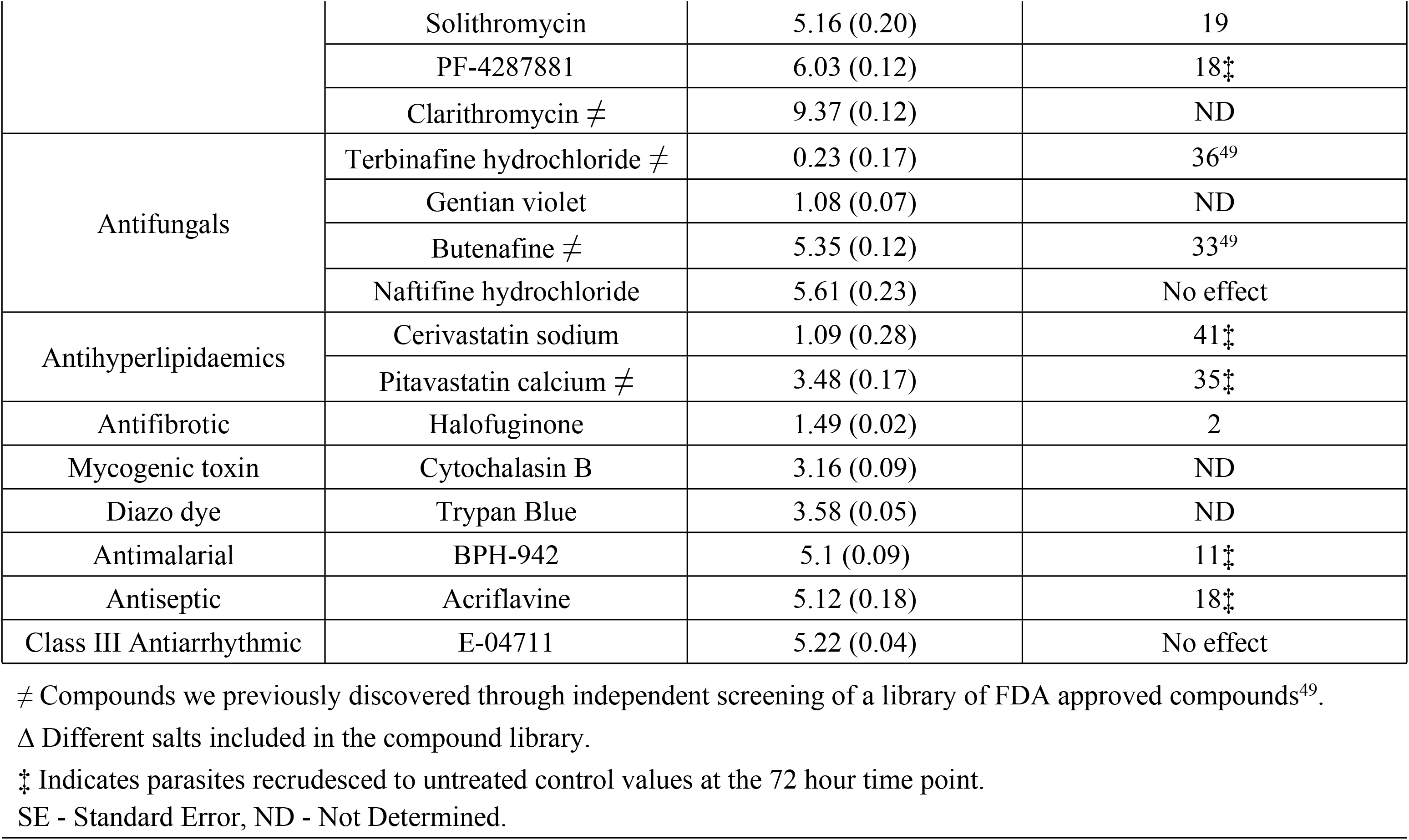
*Naegleria fowleri* active compounds identified through dose-response (N=2)

**Table 2:**
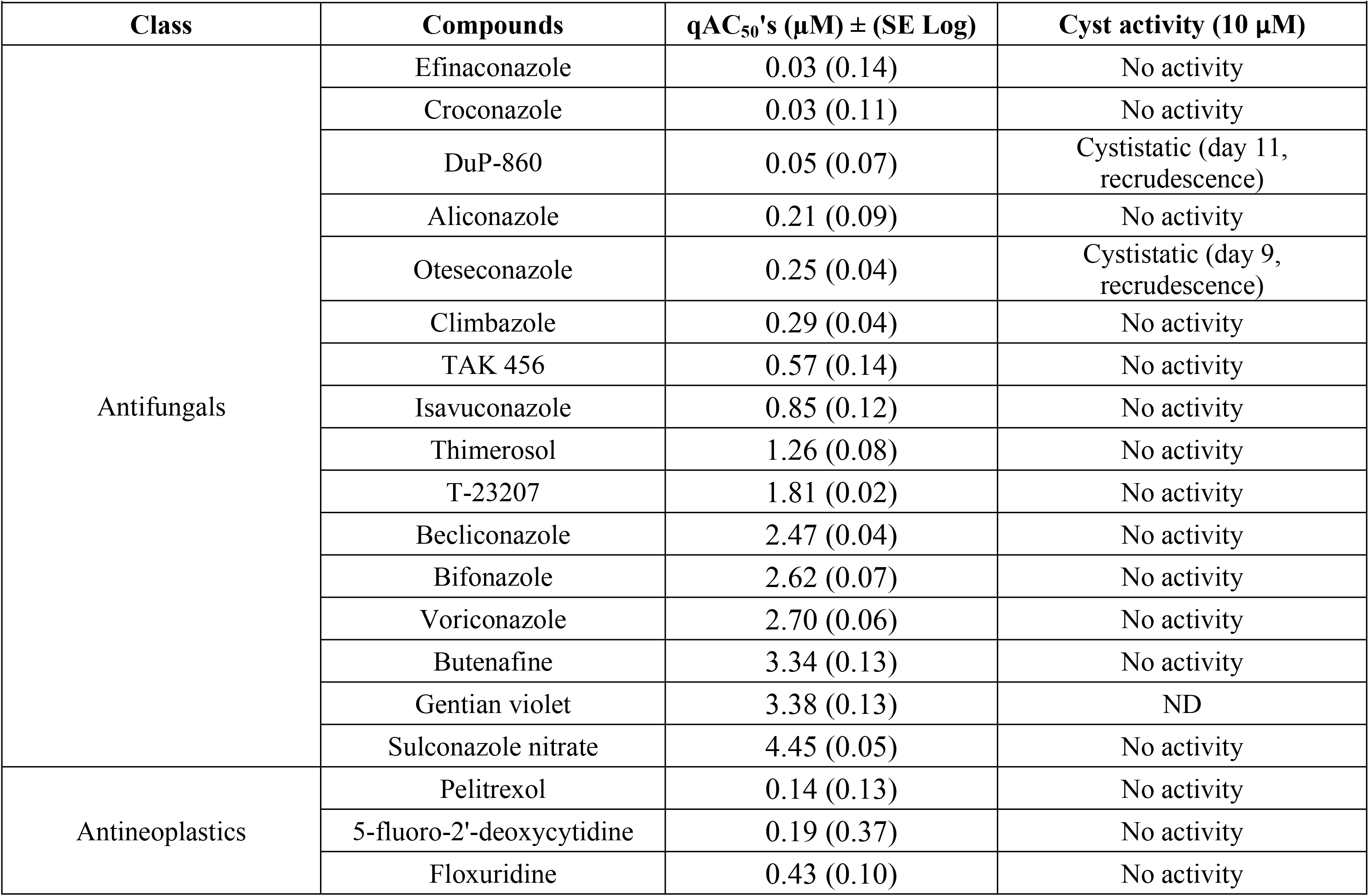
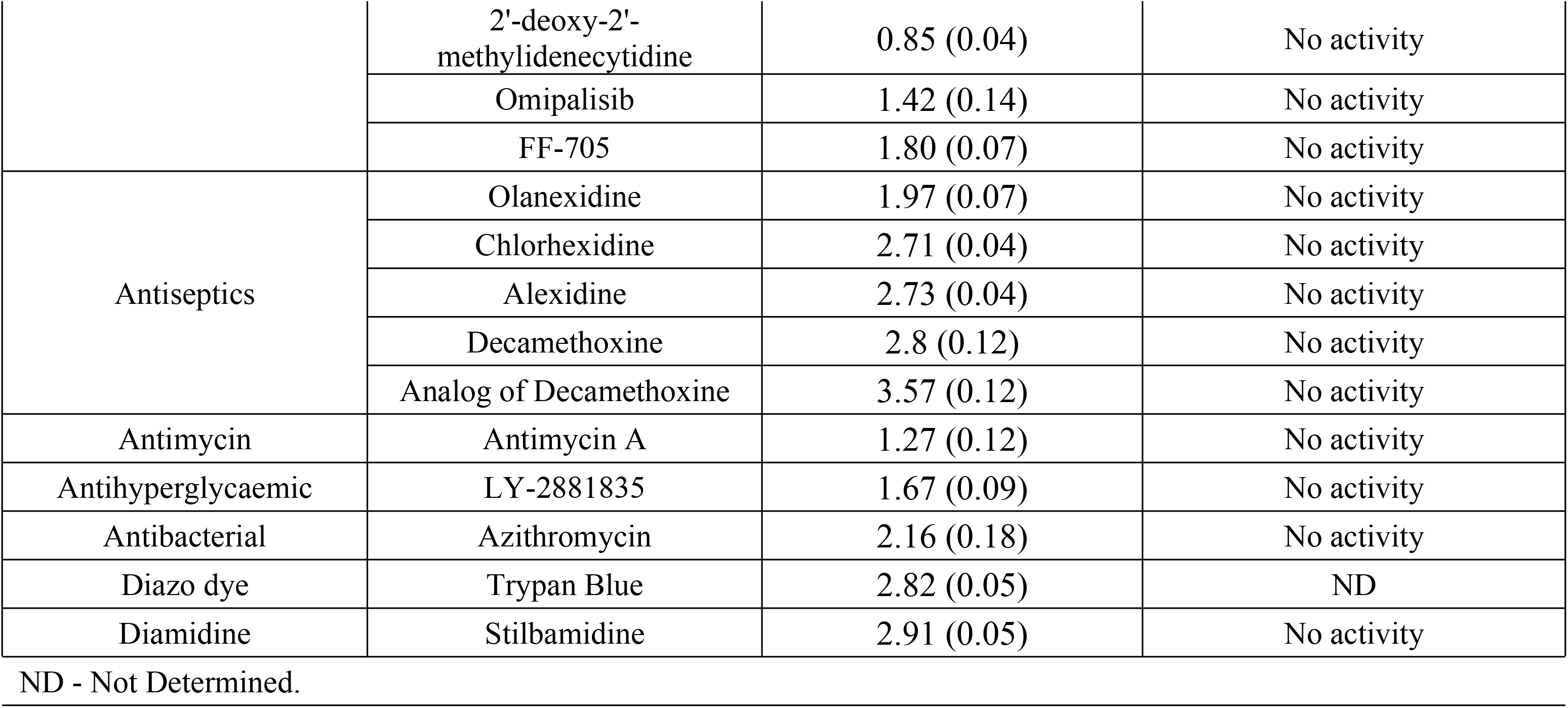
*Acanthamoeba castellanii* active compounds identified through dose-response (N=2)

**Table 3:**
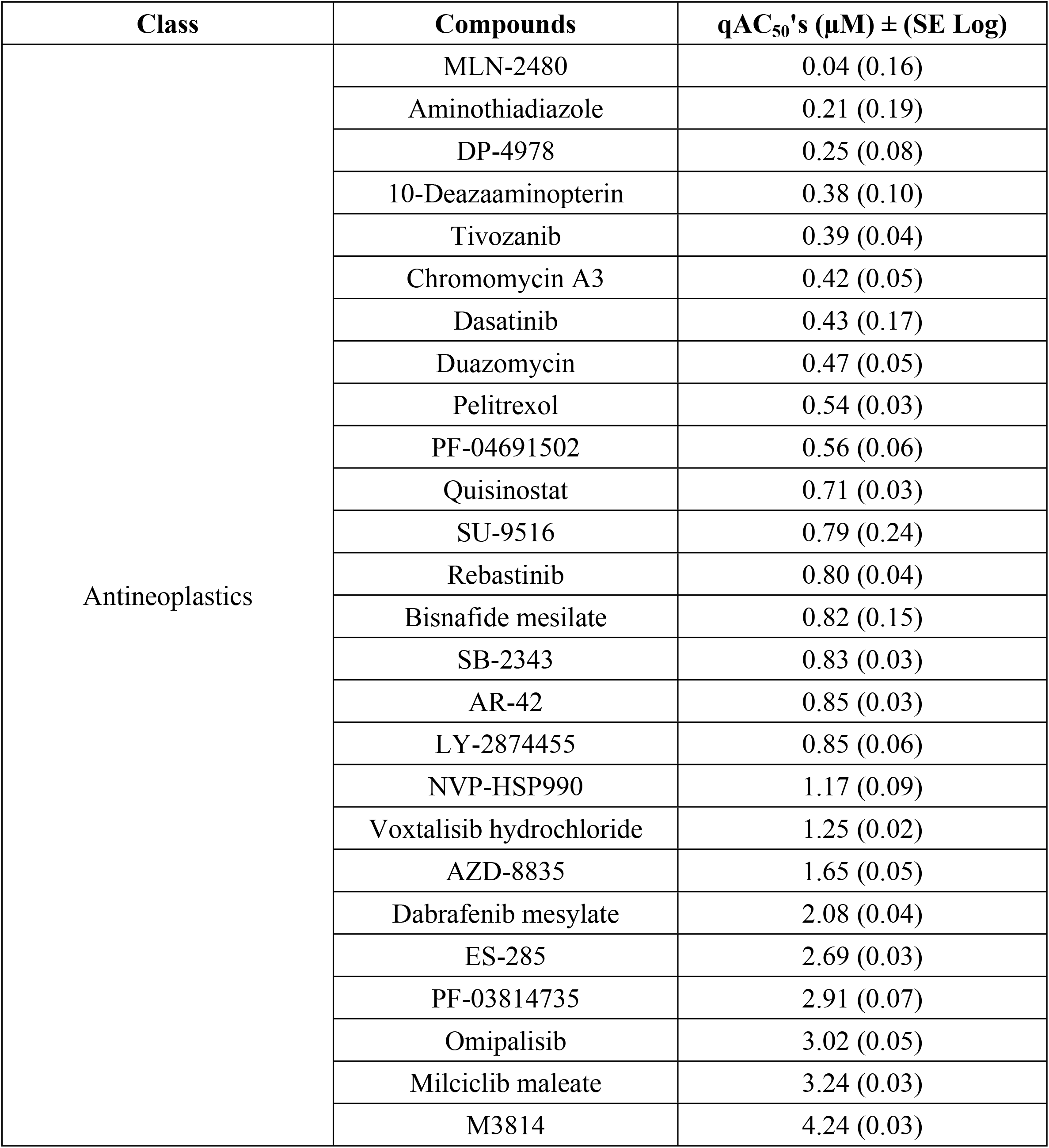
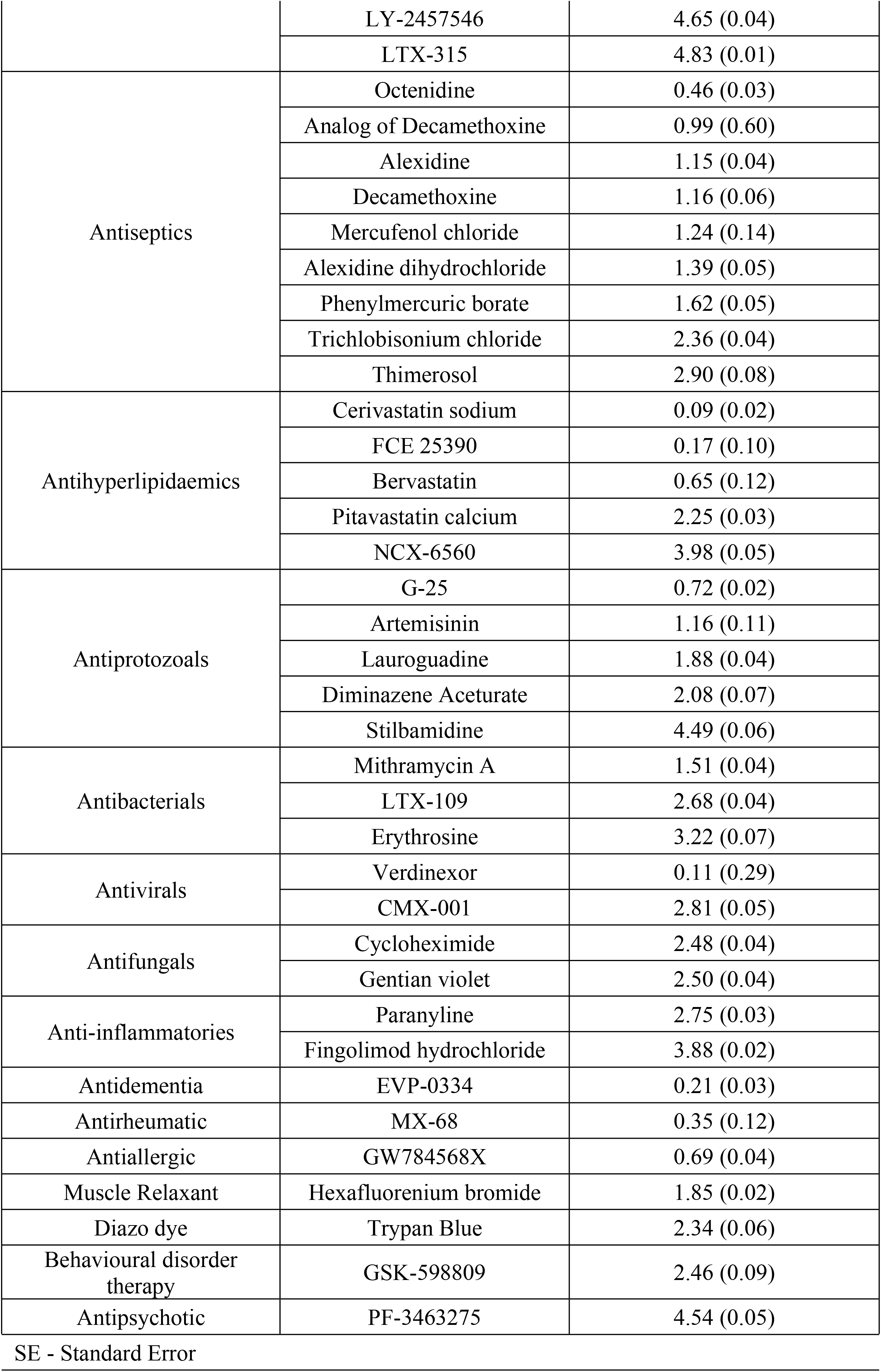
*Balamuthia mandrillaris* active compounds identified through dose-response (N=2)

### Rate of action studies

To assess the rate of inhibition of compounds on *N. fowleri*, we identified the 50% inhibitory concentration (IC_50_) for all of the confirmed hits from the single point assays, then exposed *N. fowleri* trophozoites to the IC_50_ (Fig. 2; Sup. Fig. 2) or 5x-IC_50_ (Sup. Fig. 3). Using the live-cell compatible RealTime-Glo viability reagent, luminescence was monitored kinetically every hour for a period of 72 hours and the rate of inhibition was analyzed and interpreted by calculating the area under the curve over time. At the IC_50_ concentration we identified 6 compounds having the most rapid onset of action for *N. fowleri in vitro* within 12 hours of exposure (halofuginone, NVP-HSP990, fumagillin, bardoxolone, belaronib, and BPH-942) and identified a further 13 compounds displaying inhibition within 24 hours of exposure (solithromycin, nitracrine, quisinostat, pabinostat, pracinostat, dacinostat;LAQ-824, fimepinostat, sanguinarium, radicicol, acriflavine, REP3132, BC-3205 and PF-4287881) compared to untreated *N. fowleri* control (Table 1).

**Figure 2.**
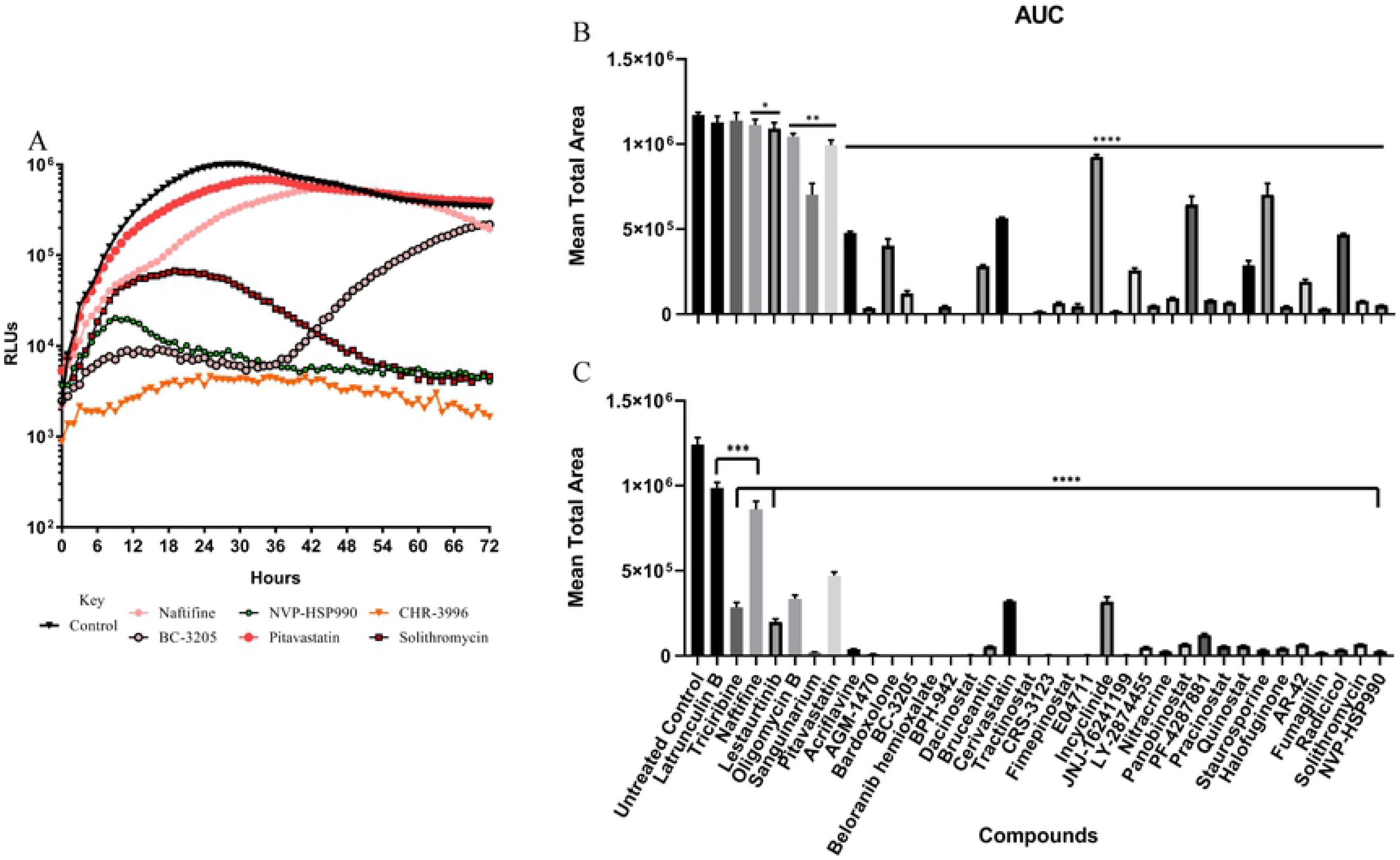
Rate of inhibition example curves and AUC analyses of the IC_50_ and 5x-IC_50_. A) Rate of inhibition at the 50% inhibitory concentration (IC_50_) for select drugs identified through screening the Calibr ReFRAME library. Data are from a single experiment with 3 replicates but are representative of independent biological repeats (N = 2). Black triangles represent untreated and normal *Naegleria* growth. RLUs, relative luminescence units. Subset of compounds from Sup. Fig. 2. B) <u>A</u>rea <u>u</u>nder the <u>c</u>urve (AUC) analysis of the IC_50_ and C) 5x-IC_50_. Multiple T tests, One unpaired T test, False Discovery Rate (FDR = 1 %) using method: Two-stage step-up method of Benjamini, Kreiger and Yekutieli was used to identify significance. Significance is recorded as *P*= < 0.05 (*), 0.0021 (**), 0.0002 (***), < 0.0001 (****), > 0.05 (NS – non-significant).

### *Acanthamoeba* cysticidal studies

After the discovery of 32 inhibitors of *Acanthamoeba* from screening the ReFRAME library (Table 2) we further assayed all of these bioactive inhibitors by determining the minimum cysticidal concentration (MCC) against fully mature cysts. We used the presence of trophozoites at 120 hours (5 days) as an indicator of cyst viability, since this is the typical time needed for strain ATCC 50370 to recrudesce and become confluent under identical culture conditions. Unfortunately, none of the compounds tested, at 10 μM, had cysticidal activity with a 24 hour exposure to inhibitors. DUP-860 and oteseconazole did display cystistatic activity with fewer trophozoites visualized after exposure to 10 μM of drug. DUP-860 treated cysts taken 4 extra days (day 9) and oteseconazole taken 6 extra days (day 11) before the trophozoites were confluent from exposure (Table 2).

### Comparison of activity of repurposed drugs for pathogenic free-living amoebae

A desired result of drug repurposing for pathogenic free-living amoebae would be the discovery of a drug(s) that significantly inhibit growth or all three major pathogens, *B. mandrillaris*, *N. fowleri*, and *A. castellanii*. Therefore, we compared the trophocidal activity of the confirmed hits and found only three compounds that inhibited all three amoebae: omipalisib, gentian violet, and trypan blue (Figure 3). Unfortunately, two of these are dyes and omipalisib was only a low micromolar inhibitor of the three pathogenic amoebae. More promising was the observation that quisinostat and AR-42, the most potent drugs identified in the screen, they inhibited both *N. fowleri* and *B. mandrillaris* with qAC_50_s < 1 μM. Similarly, pelitrexol was potently active against *A. castellanii* and *B. mandrillaris*, giving hope that at least a few of the most potent repurposing drug candidates have potential for the treatment of diseases caused by two different genus of amoebae.

**Figure 3.**
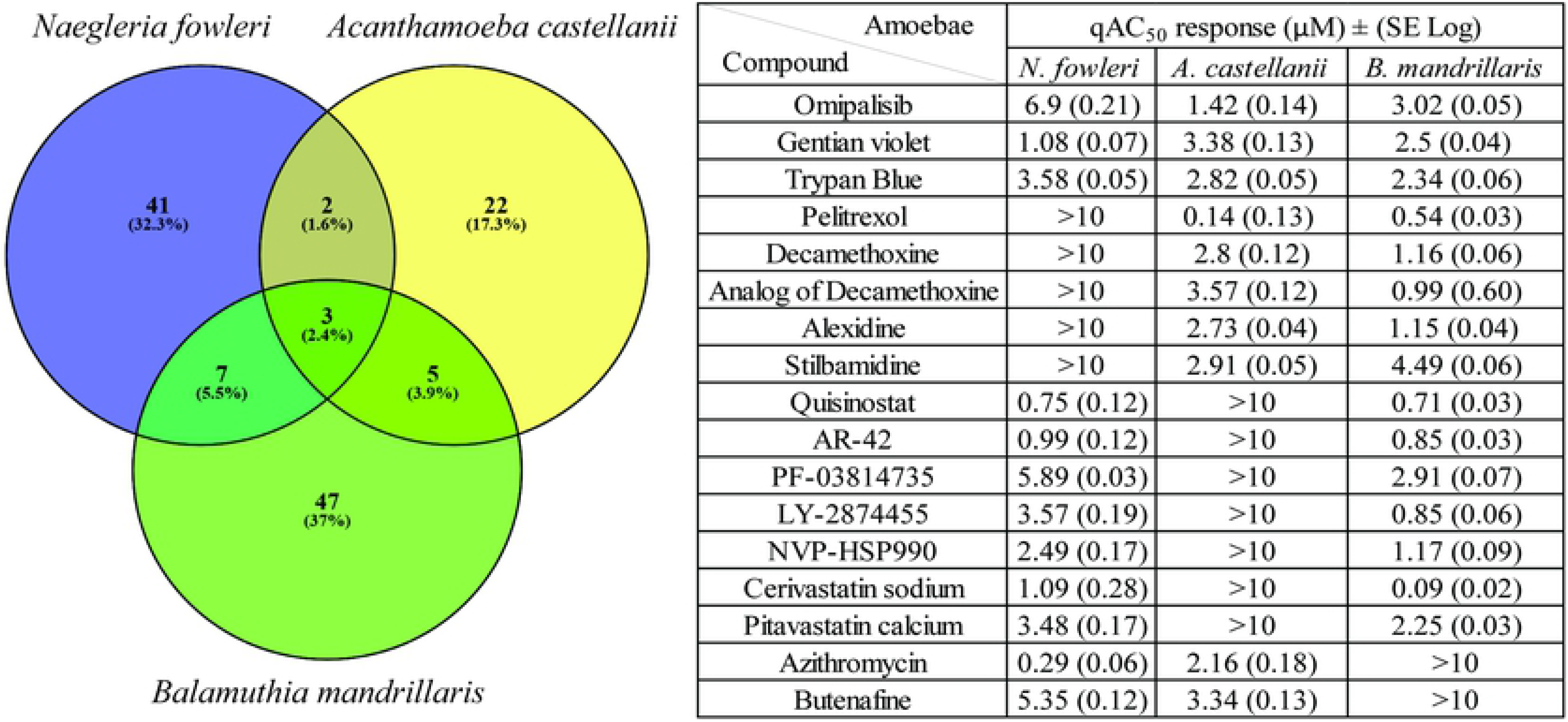
Comparison of all hits shared between the amoebae. Venn diagram illustrating the amount of individual and shared hits found between species of pathogenic amoebae. The table compares the q*AC*_50_ μM response with (SE Log).

## Discussion

To discover new drugs for repurposing as treatments for pathogenic FLA, we conducted phenotypic screening of one of the world’s largest drug repurposing libraries, the Calibr ReFRAME library. This represents the largest screening effort to date for discovery of new drugs against pathogenic *Naegleria fowleri*, *Acanthamoeba castellanii* and *Balamuthia mandrillaris*. Previously the largest drug screening efforts have included non-pathogenic amoebae as model organisms. The libraries screened include the Iconix Biosciences Inc library, Asinex compound library and a ChemDiv library on non-pathogenic *Naegleria gruberi*^44^. More recently the MMV Pathogen Box was screened against non-pathogenic *Acanthamoeba castellanii* Neff^45^. These studies extrapolated translational activity of compounds from non-pathogenic to pathogenic amoebae despite several reports of drug variation responses from different species and pathogenicity within the *Acanthamoeba* genus^46–48^. For example, we previously identified 39 agents active against pathogenic *N. fowleri* from screening the U.S. FDA approved drug library, Medicines for Malaria Venture (MMV) Pathogen Box and MMV Malaria Box^49^. The FDA drug library hits were counter screened against non-pathogenic *Naegleria lovaniensis* and we found only 2% of the active hits actually reconfirmed (unpublished data). This highlights why the discovery of new chemical scaffolds should be identified on pathogenic FLA, and further screened against multiple pathogenic species or strains for confirmation of future utility. High-throughput phenotypic cell screening provides a rapid way of robustly screening chemically diverse libraries in a cost-effective manner to identify new scaffolds or active molecules previously unknown to have any potency against FLA. Furthermore, this is the first description of a high-throughput (384 well plates) luminescence-based drug screening methodology developed for *Acanthamoeba* species.

In this study we identified 58 potent nanomolar inhibitors (19 against *N. fowleri*, 12 against *A. castellanii* and 27 against *mandrillaris*) and 90 micromolar inhibitors (34 against *N. fowleri*, 20 against *A. castellanii* and 36 against *B. mandrillaris*); 113 of these inhibitors have never been reported to have activity against these pathogenic FLA. We identified the same 8 agents from the ReFRAME library as from our previous screening of the FDA approved drug library against *N. fowleri* (azithromycin, erythromycin, roxithromycin, clarithromycin, valnemulin, terbinafine hydrochloride, butenafine and pitavastatin)^49^. This exemplifies the robustness of our high-throughput screening assays as well as true hit identification and verification.

As PAM is a disease with rapid onset and fulminant disease manifestations, an essential criterion for our target candidate profile for the development of candidate drugs against *N. fowleri* is that the drug needs to inhibit amoeba growth quickly, preferably within 24 hours of treatment. We previously reported that while azithromycin is one of the most potent inhibitors of *N. fowleri* (IC_50_ = 20 nM), it has a lag phase of ~30 hours before inhibiting growth. The compounds discovered in this study all inhibited *N. fowleri in vitro* quicker than the standard drug regimen currently used^49^. Importantly, here we describe how fluconazole and miltefosine, drugs that are currently included in the current treatment regimen, were firstly not classified as potent inhibitors within a standard *in vitro* drug susceptibility screening assays and secondly, they were not effective in our rate of action assay at the respective IC_50_. Many of the compounds discovered through screening the complete Calibr ReFRAME library inhibited the growth of *N. fowleri in vitro* quicker than the entire standard drug regimen currently recommended for the treatment of PAM^49^. We discovered 19 compounds that had a rapid onset of action and significant inhibition of *N. fowleri* within 24 hours of exposure (halofuginone, NVP-HSP990, fumagillin, bardoxolone, belaronib, BPH-942, solithromycin, nitracrine, quisinostat, pabinostat, pracinostat, dacinostat, fimepinostat, sanguinarium, radicicol, acriflavine, REP3132, BC-3205 and PF-4287881) compared to untreated *N. fowleri* control (Table 1; Sup. Fig. 2). These agents should be investigated further for in vivo efficacy and could be considered for off label indications for future treatment regimens for *Naegleria* chemotherapy.

As previously noted, clinical cases of *Acanthamoeba* have documented mixed populations of cysts and trophozoite stages in tissues during infection. Siddiqui *et al.*, reported that even with extended treatment (over 1 year) for AK, 10% of cases can still reoccur, indicating that the treatment regimen(s) failed to inactivate or kill the dormant cyst stage^50^. This is a significant problem for patients with extended treatments that can result in acute or chronic toxicity and the potential development of drug resistance to treatments. We discovered 32 compounds that had efficacy against *A. castellanii* trophozoites, 2 of which previously exhibited known cysticidal activity (chlorhexidine and alexidine) at concentrations greater than we tested at 10 μM. DuP-860 and oteseconazole displayed cystistatic activity at the highest tested concentration of 10 μM (Table 2). Cysticidal activity is one of the most important criterions for the development of candidate drugs against *Acanthamoeba* and *Balamuthia* species, due to the potential relapse of recurring infections from the protective dormant cyst stage. Unfortunately, new methods are needed to assess efficacy of drugs against cysts of *Balamuthia*.

From our initial drug susceptibility screening against *B. mandrillaris* we discovered 63 compounds that had *AC*_50_ <5 μM (Table 3). Currently the most potent compound described against *Balamuthia* is 8-Hydroxy-5-nitroquinoline (nitroxolone), discovered having IC_50_ activity of 2.84 μM^51^. In this study we identified 51 compounds more potent than nitroxolone, with activity ranging from 40 nM to 2.81 μM. In fact, this is the first description of compounds that possess nanomolar activity against *B. mandrillaris*. We confirmed anti-Balamuthia activity of 11 previously published compounds (alexidine^51^, alexidine dihydrochloride^51^, cerivastatin sodium^51^, cycloheximide^51^, dasatinib^51^, erythrosine^51^, gentian violet^51^, pitavastatin^51^, stilbamidine^51^, artemisinin^52,51^, and diminazene aceturate^53,51^, whereas the other 52 compounds we discovered are novel.

From assessing broad spectrum amoebae activity in this screen, we identified only one compound, omipalisib, and two Diazo dyes (gentian violet and trypan blue) to be active against all amoebae (Fig. 3). Omipalisib has been described as a phosphatidylinositol 3-kinase (PI3K) inhibitor, and mechanistic target of rapamycin (mTOR), We did not follow up with this agent as it only had moderate amoebae efficacy (>1 μM on all three amoebae) and is considered to be a broadly cytotoxic agent. This further demonstrates the probability that a single drug is unlikely to be potent enough against all three pathogenic FLA and that genus specific drugs may be required. In addition, rapid and specific diagnosis is required to administer an effective specific treatment tailored to the amoebae causing the infection.

Structure-based drug design as a strategy is limited, since only a handful of proteins have been crystallized and structurally investigated as potential therapeutic targets. These include genes involved in shikimate, histidine, and steroid biosynthesis, and more recently, glucose metabolism^48,54–58^. As a eukaryotic organism, these amoebae share common functional pathways with humans in which unwanted side effects can result due to the required effective concentrations needed from the current treatments used (e.g. hepatotoxicity from the use of miltefosine, nephrotoxicity due to amphotericin B use or hepatotoxicity, nephrotoxicity, dysglycemia, hyperkalemia and hyperamylasemia from the prolonged use of pentamidine^59–64^). This indicates there is a further need for studying the biology and pathogenicity of these amoebae to identify unique targets absent from the hosts.

Our repurposing efforts yielded a total of 27 potential druggable targets against *N. fowleri*, 19 against *A. castellanii*, and 44 against *B. mandrillaris*; 10 of these targets were found to be duplicated and highly conserved targets of the multiple hits identified throughout *N. fowleri*’s reconfirmed hit list (Sup. Table 1), 4 targets were conserved throughout *A. castellanii*’s reconfirmed hit list (Sup. Table 2), and 14 targets were conserved throughout *B. mandrillaris*’s reconfirmed hit list (Sup. Table 3). All of the targets identified through the current chemical inference have been requested for structural determination from Seattle Structural Genomics Center for Infectious Diseases (SSGCID)^62^.

In summary, this report describes the development of a high-throughput drug susceptibility screening methodology for *Acanthamoeba* species, discovery of new active agents that could be repurposed for future chemotherapies against pathogenic FLA infections, the identification of potential drug targets that can be used as tools for further investigation through target based therapies or structure based drug design (SBDD) and we describe the identification of novel compounds that inhibit *N. fowleri* quicker than any of the standard drug regimen currently used.

Given the limited availability of effective drugs against pathogenic FLA, our results offer a plethora of active drugs against *N. fowleri*, *A. castellanii* and *B. mandrillaris* that warrant further exploration for treatment of diseases caused by pathogenic FLA.

## Author contributions

C.A.R., B.L.C., E.C., M.V.H., and D.E.K designed the experiments. C.A.R and B.L.C performed the experiments. C.A.R., D.E.K., and M.V.H., analyzed the data. E.C. prepared the drug library. C.A.R. and D.E.K drafted the manuscript. All authors participated in critical evaluation and revision of the manuscript before submission.

## Acknowledgements

This work was supported by the Georgia Research Alliance and the Bill & Melinda Gates Foundation.

The authors would like to thank Dr.’s Case W. McNamara, Arnab K. Chatterjee, and Peter G. Schultz for the curation of the Calibr ReFRAME library as well as all of the staff involved at Calibr, a division of The Scripps Research Institute, for the assembly and distribution of this unique repurposing compound library.

